# Successful field performance in dry-warm environments of soybean expressing the sunflower transcription factor HaHB4

**DOI:** 10.1101/2019.12.21.885798

**Authors:** KF Ribichich, M Chiozza, S Ávalos-Britez, JV Cabello, AL Arce, G Watson, C Arias, M Portapila, F Trucco, ME Otegui, RL Chan

**Author notes:** these authors equally contributed to this work (KF Ribichich, M Chiozza). these authors equally contributed to this work (S Ávalos-Britez, JV Cabello, AL Arce). these authors are corresponding authors (ME Otegui, RL Chan). Corresponding authors: María Elena Otegui, Raquel Lía Chan. **Authors’ mail addresses:** Karina Ribichich, Mariana Chiozza, Selva Álvarez Brítez, Julieta Cabello, Agustín Arce, Claudia Arias, Margarita Portapila, Gerónimo Watson, Federico Trucco.

## Abstract

Soybean yield is limited primarily by abiotic constraints. No transgenic soybean with improved abiotic-stress tolerance is available in the market. We transformed soybean plants with genetic constructs able to express the sunflower transcription factor HaHB4, which confers drought tolerance to Arabidopsis and wheat plants. One line (b10H) carrying the sunflower promoter was chosen among three independent lines because it exhibited the best performance in seed yield (SY). Such line was evaluated in the greenhouse and in twenty-seven field trials developed in different environments of Argentina. In greenhouse experiments, transgenic plants showed increased SY under stress conditions together with wider epycotyl diameter, enlarged xylem area and enhanced water use efficiency than controls. They also exhibited enhanced SY in warm-dry field conditions. This response was accompanied by the increased in seed number that was not compensated by the decreased in individual seed weight. The transcriptome analysis of plants from a field trial with maximum SY difference between genotypes indicated an induction of genes encoding redox and heat shock proteins in b10H. Collectively, our results indicate that soybeans transformed with *HaHB4* are expected to have reduced SY penalization when cropped in warm-dry conditions, which constitute the best target environments for this technology.

**Highlight:** Soybean transformed with the sunflower gene encoding the transcription factor HaHB4 was evaluated in greenhouse and field trials. Transgenic plants significantly outyielded controls in drought-warm environments due to, at least in part, increased seed number, xylem area, and water use efficiency as well as to the induction of genes encoding redox and heat shock proteins.

## Introduction

Soybean (*Glycine max* L. Merr.) is one of the most important crops in the world, including a wide range of uses. Many countries adopted biotech soybeans in more than 90 percent (http://www.isaaa.org/). However, biotic and abiotic constraints still limit seed yield (SY) and seed quality of this species (Hartman *et al.*, 2011).

Approved and commercialized genetically modified (GMO) soybean involved resistance to herbicide or herbivory attack, whereas GMOs with increased abiotic stress tolerance remain absent from the worldwide market until now. Regarding other species and regions/countries, increased seed yield (SY) of genetically modified (GM), drought-tolerant maize grown under water-limited conditions was described by Castiglioni *et al.* (2008). Such maize plants express the bacterial RNA chaperons CspB and CspA, which generated drought tolerance as well as cold and heat tolerance (Liang, 2017).

Regarding transgenic soybean, besides the works devoted to glyphosate technology, scientific literature considering other traits is scarce, and almost absent when field trials are requested. A few manuscripts reported evaluation of transgenic soybean, mostly in greenhouse or growth chamber conditions. Such is the case of soybean plants overexpressing *GmSYP24*, a dehydration-responsive gene showing insensitivity to osmotic/drought and high salinity stresses via stomata closure involving an ABA signal pathway in greenhouse assessments (Chen *et al.*, 2019). Another work described the overexpression of the b-Zip transcription factor (TF) GmFDL19 showing early flowering and enhanced tolerance to drought and salt stress; however, SY in stressing or standard growth conditions was not informed (Li *et al.*, 2017). Similarly, a soybean MYB encoding gene was overexpressed and tested in soybean plants. Transgenic soybean plants carrying an extra copy of *GmMYB84* (a R2R3-MYB TF) exhibited a higher survival rate after severe drought when they were tested in controlled conditions in a culture chamber (Wang *et al.*, 2017).

Water deficit is the most important factor affecting the crop SY worldwide. Water scarcity for agriculture increases the production costs and determines the need of improving the resource use efficiency across a broad range of permanent as well as transient drought-prone regions of the world.

Drought tolerance without yield penalties is a desirable trait but difficult to achieve. Plants evolved to survive under water deficit conditions displaying physiological changes which especially include stomata closure. Most of the genes, positively involved in drought response and tested both in model plants and crops, induce stomata closure and hence, increase plant survival but reduce biomass and seed production under the very frequent mild stress conditions (Skirycz *et al.*, 2011). Moreover, Passioura (2012) analyzed a huge number of reports referred to drought-tolerant Arabidopsis plants and detected that the enhanced survival of a high percentage of such plants was simply explained by their reduced size and concomitant slowed water uptake than the wild type (Morran *et al.*, 2011).

TFs are particularly abundant in the plant kingdom, representing about 6% of the encoded proteins (Riechmann *et al.*, 2002). Among plant TFs, homeodomain-leucine zipper (HD-Zip) proteins are unique to plants and have been assigned roles in development associated to environmental stressing factors (Ariel *et al.*, 2007; Perotti *et al.*, 2016). Although their conserved structures and functions, HD-Zip I TFs diverged during evolution, presenting differential features when comparing plant species.

Soybean plants have 36 members of the HD-Zip I family (Belamkar *et al.*, 2014). Among them, *GmHB6, GmHB13* and *GmHB21* showed different expression levels after drought treatment in susceptible (BR 16) and tolerant (EMBRAPA 48) soybean cultivars, indicating the presence of differential regulation *cis-acting* elements. Particularly, GmHB13 was exclusively induced by water deficit in the tolerant cultivar whereas GmHB6 was only repressed in the susceptible one (Pereira *et al.*, 2011). Functional studies about these TFs are not available to date in the scientific literature but their differential expression in tolerant and susceptible cultivars suggests a role in the response to drought.

Sunflower belongs to the Asteraceae clade of Angiosperms and have several divergent HD-Zip I members (Arce *et al.*, 2011). Among them, HaHB4 (*<u>H</u>elianthus <u>a</u>nnuus <u>H</u>omeo<u>B</u>ox* 4) has been deeply characterized. This TF exhibits an abnormal short carboxy terminus compared with Arabidopsis members, and its expression is highly induced by several environmental factors (drought, salinity, darkness) and hormones (ethylene, ABA, jasmonic acid) (Gago *et al.*, 2003; Manavella *et al.*, 2006, Manavella *et al.*, 2008a, 2008b, 2008c). Arabidopsis plants expressing this sunflower TF, either under a constitutive or inducible promoter (Cabello *et al.*, 2007), exhibited enhanced tolerance to water deficit.

Recently, it was reported that HaHB4 was able to confer drought tolerance to wheat plants tested in greenhouses and in 37 field trials (González *et al.*, 2019). It was proposed as part of a potential molecular mechanism that this TF could interact with endogenous members of the same family by dimerization or by protein-DNA interactions (Gonzalez *et al.*, 2019).

In this work we show that HaHB4 was able to confer drought tolerance and increased SY to soybean plants tested in the field, particularly in warm dry environments. Moreover, we show here that the improved performance of HaHB4 plants is strongly related to enhanced water uptake, biomass production and water use efficiency as well as to differential plant architecture. Transcriptome analyses performed with field-harvested samples indicated that redox and heat shock proteins encoding genes are induced in the transgenic genotype. This work constitutes a multidisciplinary approach contributing to understand abiotic stress tolerance mechanisms displayed in soybean by the introduction of the sunflower transcription factor HaHB4.

## Materials and Methods

### Genetic constructs

The open reading frame of cDNA encoding full-length *HaHB4* cloned in the *BamHI/SacI* sites of *pBluescript SK-* (Stratagene, Upsala, Sweden) was used as template in a PCR reaction with oligonucleotides H4-F (5’ ATGTCTCTTCAACAAGTAACAACCACCAGG-3’) and Transf2 (GCCGAGCTCTTAGAACTCCCACCACTTTTG-3’), which included initiation and stop codons. The PCR amplification product was cloned into a pGEM®-T-Easy vector (Promega, Madison, Wisconsin) and named pHaHB4.2. Then, the cDNA was cloned in expression cassettes bearing two different promoters: (a) the constitutive *35S CaMV* promoter (*35S:HaHB4.2*) and (b) the inducible *HaHB4* promoter. Both cassettes were subcloned in a vector carrying the bar gene and the NOS termination sequence (Chan and González, 2013). Clones were obtained in *Escherichia coli* and then *Agrobacterium tumefaciens* (strain EHA101) was transformed. The sequences were checked (Macrogen Korea) and, as previously described, a few mutations detected (Chan and González, 2012; González *et al.*, 2019). Transactivation activity and other characteristics of such point mutants were described in detail in González *et al.* (2019).

### Plant transformation and selection of transgenic events

Soybean transgenic events were generated using an *Agrobacterium*–mediated protocol and the cultivar Williams 82 (hereafter W82) according to the methods described by Hofgen *et al.* (1998). Transgenic events were selected using ammonium glufosinate. T_1_ seeds were obtained for 35 independent events.

Multiplication of the transformed cells was conducted in a greenhouse. T_1_ individuals derived from each event were sampled for a segregation test by PCR determination. Lines derived from selfings of individuals from selected events (3:1 segregation in T_1_) were sowed and analyzed by PCR to identify homozygous lines, as indicated by the absence of negative segregants among the sampled progeny (at least 5 individuals sampled per line). Seed augmentation (T3 seed) of single-copy homozygous was conducted in a greenhouse.

### Field trials for event selection and cultivars testing

The experimental network for the evaluation of HaHB4 effects in soybeans included 30 field experiments conducted across a wide environmental range through several growing seasons (6) and sites (16). Experiments were organized in three groups. Group 1 corresponded to the evaluation of 3 transgenic (TG) events (a11H, a5H, and b10H) respect to the wild type (WT) parental cv. W82 and included 17 experiments developed between 2009-2010 and 2012-2013. Some of these experiments were also included as part of Group 2 (described next). General growing conditions (i.e. cumulative incident solar radiation, mean temperatures, rainfall, and potential evapotranspiration) along the cycle experienced by crops in 14 of these experiments are described in Supplementary Table 1. Group 2 corresponded to the analysis of the best performing event (b10H) respect to the WT parental W82 for the detection of genotype (G) per environment (E) interactions (G×E) and included 27 experiments carried out between 2009-2010 and 2018-2019. For this group, an environmental index (EI) was computed as the average seed yield (SY) or SY component (seed numbers; individual seed weight) of all evaluated genotypes in a given environment. Each trait of b10H as well as of W82 in each environment was regressed respect to the corresponding EI. Growing conditions of all experiments in Group 2 are described in Supplementary Table 1. Rainfall data were obtained *in situ*, whereas other weather records were obtained from the nearest weather station (http://siga2.inta.gov.ar). Water balance for different growth periods and for the whole cycle was obtained as the difference between potential evapotranspiration (PET, in mm) and water supplied by rainfall (Rain, in mm) plus irrigation (Ir, in mm). The relative water balance (RWB) was computed as in Eq. 1.

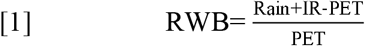

Group 3 focused on the detection of differences in the physiological determination of SY between bH10 and W82, which was performed in 4 of the 27 experiments mentioned for Group 2 (those developed during 2017-2018 and 2018-2019).

Some experiments were carried out in the same year × site combination but using different sowing dates (e.g. Aranguren-2013, Carmen de Areco, Roldán and San Agustín sites), water regimes (e.g. Liborio Luna, and Pergamino) or phosphorus fertilizer rate (e.g. Aranguren-2014). Seed yield of all cvs. was always assessed in a randomized complete block design with at least 3 replicates and plots of at least 10.4 m^2^ (2.08 m width × 5 m length). Plots were machine-sown in all experiments except those performed at Pergamino and IAL-Santa Fe (2017-2018), which were hand-sown. Sowing date took place between 7-Nov and 14-Jan, harvest occurred between 28-Mar and 9-May, and the stand density ranged between 30-40 plants m^−2^.

At the 2017-2018 water-deficit experiment of Pergamino, rainfall water was excluded from plots by means of removable shelters installed 23 days after sowing (i.e., before the start of R1 on 39 d after sowing; Fehr and Caviness, 1977) and removed 91 d after sowing (ca. R6). In this experiment, soil water content was surveyed from sowing+23 days up to R7 (on 110 d after sowing) by means of volumetric (0-30 cm) as well as neutron probe (Troxler 3400, NC) measurements (30-185 cm). The difference between successive soil water measurements plus the amount of Rain+Ir water added to each plot allowed estimation of crop water use (ET_C_: crop evapotranspiration, in mm) during the mentioned period. All experiments were kept free of weeds, insects and diseases by means of the necessary recommended controls.

### Crop and plant phenotyping in field experiments

Days to R1 and R7 were assessed in 11 experiments of Group 1 for the analysis of event effects on phenology. In all experiments SY was obtained at R8 by machine (all experiments except those performed at Pergamino and IAL-Santa Fe sites) or hand harvesting (at Pergamino and IAL-Santa Fe sites) all plants present in at least 1 m^2^ of a central row of each plot, which were threshed for seed recovery. Seeds were cleaned and weighed, and seed weight corrected for estimation of SY (in g m^−2^) on a 13% wet basis. Relative SY of each experiment was computed as in Eq. 2.

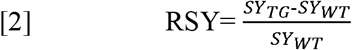

where SY_TG_ and SY_WT_ represent SY of the transgenic b10H and of the wildtype parental W82, respectively.

The number of seeds (seed numbers) and individual seed weight were assessed in 17 experiments of Group 1, 23 experiments of Group 2 and all experiments of Group 3. For this purpose, at least 3 samples of 100 seeds each were taken from the seed bulk and weighed; the obtained values were averaged for estimation of individual seed weight (in mg). Seed number was computed as the ratio between SY and individual seed weight and expressed on a per m^2^ basis.

Total aerial crop biomass per m^2^ (BIOM/m^2^, in g m^−2^) as well as pod biomass per m^2^ (POD_B_/m^2^, in g m^−2^) and pod numbers per m^2^ (POD_N_/m^2^) were surveyed at physiological maturity (R7) at the Pergamino and IAL-Santa Fe sites. For this purpose, plants present in 0.52 m^2^ were collected from a central row in each plot and dried at 60°C until constant weight. The number of pods with at least one developed seed was counted on these plants. Biomass partitioning to reproductive organs at R7 was estimated as (i) the ratio between POD_B_/m^2^ and BIOM/m^2^, described as biomass partitioning index to pods (BPI_P_), and (ii) the ratio between SY and BIOM/m^2^, described as harvest index. At Pergamino, water use efficiency (WUE) based on crop evapotranspiration (ET_C_) was computed for biomass (WUE_B,ETc_) as well as for seed (WUE_SD,ETc_) production. The former was obtained as the ratio between BIOM/m^2^ and ET_C_ whereas the latter was obtained as the ratio between SY and ETC.

At the IAL site, plants at reproductive stage (R1, R3 and R5) were evaluated including light interception measured with a ceptometer (Cavadevices, Argentina) as described in Maddonni and Otegui (1996). Midpoint internode diameters (hypocotyls and epicotyls) were measured on 3 plants at V2 stage and branches per plant on three plants plot^−1^ at R5. Stem sections were collected from the same region and treated as described below. Relative xylem area was estimated as xylem area/total stem area measured with Image J (Rasband, 1997-2018).

### Greenhouse growth conditions and plant phenotyping

A greenhouse experiment was performed at the IAL site. Seeds of the WT (W82) and the TG (b10H) were sown and grown in 0.5 L pots filled with white peat (klasmann-deilmann TS1) during two weeks in a culture chamber (18 h light photoperiod, 23±1°C). Then, 14 plants per genotype were individually grown each in 8 L pots filled with peat (Terrafertil Growmix Multipro): perlite (3:1) and 1.25 g L^−1^ of slow release fertilizer (Compost Expert Basacote Plus) and grown until harvest in a greenhouse under temperature and humidity monitoring. One week after placing the plants in the greenhouse, 50% of the plants were subjected to mild water stress watering the pots to 60% of field capacity up to R3 (53 days). The rest of the plants (controls) remained well-watered to 100% of field capacity during the treatment period, and pots of all plants were watered up to field capacity from R3 onwards.

Plant water uptake was estimated from the difference in pot weight between pots held at 100% and 60% of field capacity, considering negligible the weight of plants and the soil evaporation (plants at V5 and older cover the pot surface). Accumulated water uptake was computed for each plant as total water (ml) added during the stress period. Relative water uptake (ml) was estimated as the ratio between water uptake by water-deficit and control plants of each genotype.

Yield components, plant height, number of branches, internodes and pods per plant were registered at final harvest. Midpoint internode diameters (epicotyl and 1^st^, 2^nd^ and 3^rd^ internodes) were measured on 3-4 plants at V5 with water-deficit treatment (no water addition between V3 to V5) or without it. Stem sections and xylem area were estimated with Image J (Rasband, 1997-2018).

### Histological and microscopic analyses

Stem sections of 0.5-1.0 cm length were collected and fixed at room temperature for 24 h in FAA solution (3.7 % formaldehyde, 5 % acetic acid and 50 % ethylic alcohol), and then subjected to standard alcohol series dehydration and paraffin (Histoplast (Biopack^™^, Argentina) inclusion protocols (D’Ambrogio de Argueso, 1986). Transverse stem sections (10 μm thick) were obtained using a microtome (RM2125, Leica). Sections were mounted on slides coated with 50 mg/ml poly-d-Lys (Sigma Chemical Co., St. Louis, MO, US) in 10 mM Tris-HCl pH 8.0 and dried during 16 h at 37 °C. After removing the paraffin, the slices were treated with safranine-fast green staining (D’Ambrogio de Argueso, 1986), and mounted on Canadian balsam (Biopack™, Argentina) for microscopic visualization in an Eclipse E200 Microscope (Nikon, Tokyo, Japan) equipped with a Nikon Coolpix L810 camera.

### Evaluation of photosynthetic parameters in the field trial at the IAL site

Photosynthetic parameters were measured in TG (b10H) and WT (W82) soybeans during 2017-2018 at the IAL site. Measurements were made on healthy and fully expanded leaves of plants randomly chosen and during different growth stages (R3, R5 and R6). The net photosynthetic rate (P_n_) was assessed with a portable photosynthesis system (LI-COR, Lincoln, Nebraska, USA). Photosynthetically active radiation (PAR) was provided by a LED light source set to 1500 μmol m^−2^ s^−1^, air flow rate through the sample chamber was set at 500 μmol^−1^ s^−1^and CO_2_ concentration was 400 μmol mol^−1^. Air relative humidity range was 50-60 % and leaf temperature range was 25-30°C.

### Transcriptome analysis by RNA-Seq

Three leaf fragments (around 1 cm^2^ each) of 8-10 plants per plot, were collected at the IAL field assay, sown in liquid nitrogen and stored at −80°C. Samples from R5-R6 were used for RNA-Seq. Total RNA was extracted with RNAeasy (Quiagen) from pulverized samples. RNA quality and integrity were checked by absorbance (260/280 > 1.8, 260/230 > 2.0) and electrophoresis. RNA was analyzed by BGI (San Jose, USA) by sequencing 8 libraries. An average of 82,468,181 clean reads/sample with more than 95% of them with Q>20 were reported.

Raw paired-end reads were first quality trimmed with Trimmomatic (version 0.36; Bolger *et al.*, 2014) and then aligned to the Glycine max W82 genome, v4 (Schmutz *et al.*, 2010; from Phytozome V13, Goodstein *et al.*, 2012) using STAR (version 2.5.2b, Dobin *et al.*, 2014) with a maximum intron length of 1200 bp. Using samtools (version 1.8; Li *et al.*, 2009), only primary alignments with a minimum MAPQ of 3 were kept. Read quality before and after trimming was analyzed with FastQC (version 0.11.5; Andrews 2010) and, together with mapping efficiency, was summarized with MultiQC (version 1.7; Ewels *et al.*, 2016). Read counts on each gene were calculated with featureCounts (version 1.6.2; Liao *et al.*, 2014) using the gene and exon annotation from Phytozome (V13, Goodstein *et al.*, 2012). Differentially expressed genes were determined with DESeq2 (Love *et al.*, 2014; R Core Team, 2018) filtering out genes with counts below 10 in all samples. This analysis pipeline was run with the aid of the Snakemake workflow engine (Köster and Rahmann, 2012). Gene ontology analysis was performed online with agriGO (v2, Tian *et al.*, 2017).

### Statistical analyses

Differences in SY and its components between WT cv. W82 and TG events (experiments in Group 1) as well as between W82 and TG cv. b10H (experiments in Groups 2 and 3 as well as in the greenhouse) were assessed by means of analyses of variance (ANOVAs), with genotypes (G) and environments (E) as fixed factors and replicates nested within environments. A Tukey test was used for comparison of main and interaction (G×E) effects. Square root transformation was used to transform discrete variables. Other traits within a given environment were evaluated by *t* test. The relationship between variables was evaluated by correlation and regression analyses.

The photosynthetic parameters evaluation was performed using the statistical software package SSPP 20.0 (SSPP Inc., Chicago, IL, USA).

### Accession numbers

For sunflower *HaHB4*, accession numbers in EMBL, GenBank and DDBJ Nucleotide Sequence Databases are AF339748 and AF339749.

The IDs of differentially expressed soybean genes identified in the RNA-Seq are listed in Supplementary Table 2.

## Results

### A set of field trials allowed the selection of a soybean transgenic HaHB4 event

Different transgenic lines bearing either the constitutive 35S (lines called “a”) or the HaHB4 (lines called “b”) promoter were obtained and, together with the WT cv. W82, were multiplied and evaluated in field trials. After a first assessment, three independent events (a5H, a11H and b10H) bearing only one copy of the transgene were selected for further assessment.

From the experiments developed for event selection (Group 1), it could be established that the presence of HaHB4 produced (i) no effect on days to R1 (data not shown), (ii) a slight delay on days to R7 (data not shown), (iii) increased SY (Fig. 1A) due to increased seed numbers (Fig. 1B), and (iv) decreased individual seed weight (Fig. 1C). No event expressing *HaHB4* differed from the WT in days to R1 (i.e. beginning bloom). Across experiments, the WT took 42.5±9.5 days to R1, whereas the shortest event (b10H) took 42.3±9.2 days and the longest event (a5H) took 42.8±9.0 days to this stage (i.e. an average of only 0.43 d between the longest and the shortest cvs). By contrast, all events expressing *HaHB4* tended to have a delayed senescence respect to the WT, though the number of days to R7 (i.e. beginning maturity) was slightly modified among them (maximum range of 1.66 days across mean values). The difference with the WT (mean of 113.9±10.4 d to R7), therefore, was significant (*P<0.05*) only for the a5H event (mean of 115.6±10.7 d to R7), and final harvest was done on the same date for all genotypes. The trade-off between increased seed numbers and decreased seed weight was only partial for b10H (SY larger than SY of W82) and total for the other events (SY equal to SY of W82), and consequently b10H was selected for subsequent studies.

**Figure 1.**
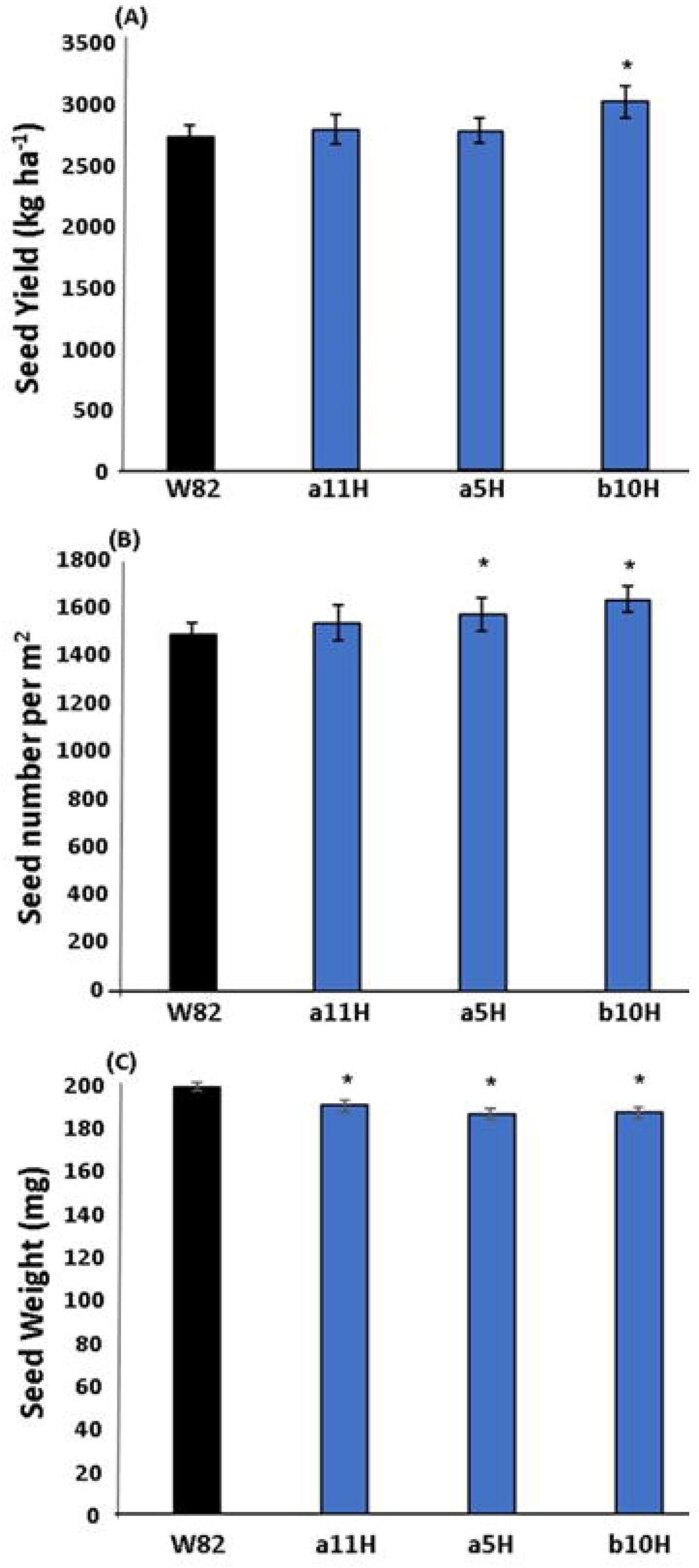
Comparative performance between transgenic events and wild-type parental W82. Figures on the left represent (A) Seed yield, (B) Seed number m^2^, and (C) Individual seed weight. Data are mean values ± SEM×2 of four soybean genotypes (the wild type W82 and three transgenic lines). The asterisk indicates significant difference (P<0.01) respect to the wild type. Figures on the right represent the comparison between the transgenic cv. b10H and the parental cv. W82 across all evaluated environments (Supplementary Table 1). The mean value of each environment corresponds to the average of all tested genotypes and is described as an environmental index (EI). Fitted models in (D) differed at P<0.05 and indicated that b10H will outyield W82 across all environments with seed yield lower than 4898 kg ha^−1^, a threshold never met in current research. No cross-over interaction was detected for models fitted to seed numbers (E) and seed weight (F). Ordinates of models fitted in (E) and (F) differed at P<0.0001.

### Transgenic soybean significantly outyields its control in field trials

From the results of 27 field experiments performed across a wide environmental range (Fig. 2A), it could be established that the TG cv. b10H significantly outyielded the parental WT cv. W82 (Fig. 2B). This advantage averaged +4.05% (range between −11% and +43%) and held across all the environmental range explored (Supplementary Fig. 1), which extended from a minimum of 1540 kg ha^−1^ to a maximum of 4540 kg ha^−1^. Models fitted to the response of each genotype to the environmental index indicated that b10H outyielded W82 across all environments with seed yield lower than 4898 kg ha^−1^. This threshold was never met among evaluated environments. The described SY advantage of b10H was supported by the larger number of seeds (mean of +10.6%, Fig. 2C), which was not compensated by the reduction registered in individual seed weight (mean of −6.5%, Fig. 2D). No cross-over interaction was detected for SY components across environmental indexes (Supplementary Fig. 1), being b10H>W82 for the main determinant of SY (i.e. seed number m^−2^, Fig.1B).

**Figure 2.**
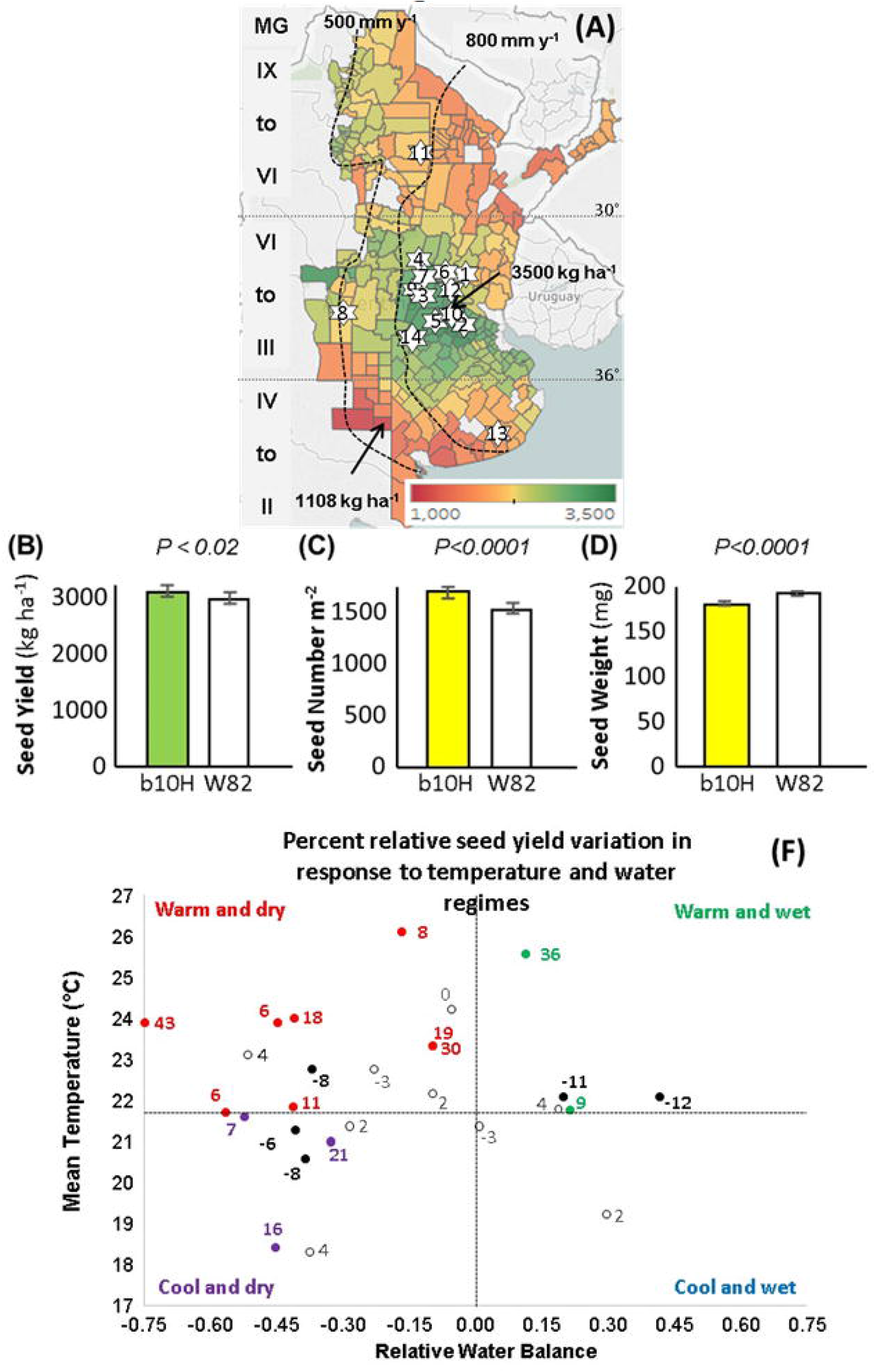
Transgenic cv b10H outyields the WT cv W82 across 27 field-based experiments, particularly in dry-warm environments. (A) Location of the 14 sites corresponding to the experimental network of Indear (details in Supplementary Table 1). The colored scale in the map represents the 22-y mean seed yield of soybeans for the period between 1996 (year of release of Roundup Ready cvs) and 2018 in Argentina. Non-colored areas correspond to counties with less than 10 records for the evaluated period, whereas arrows indicate the counties with maximum (3500 kg ha^−1^) and minimum (1108 kg ha^−1^) mean seed yields. Roman numerals on the left describe the currently recommended maturity groups (MG) across the latitude gradient. Dashed lines indicate the 500 and 800 mm year^−1^ isohyets. Mean values of seed yield (B), seed number m^−2^ (C) and individual seed weight (D) of cvs b10H and W82 across 27 experiments. Error bars represent SEM×2. (E) Relationship of mean temperature and mean relative water balance of each experiment (n = 27; Supplementary Table 1). Dashed lines represent the median mean temperature (horizontal) and the null relative water balance (vertical). The values next to each symbol represent the relative seed yield (RSY= (SYtg-SYwt)/SYwt; SYtg: seed yield of transgenic cv. b10H; SYwt: seed yield of WT parental cv. W82). Different colors represent cases with (i) RSY ≥ 0.05 (SYtg>SYwt), in bolded non-black colors that identified the environmental group represented in each quadrant, (ii) RSY ≤ −0.05 (SYwt>SYtg), in bolded black, and (iii) −0.05<RGY<0.05 (SYtg ≅ SYwt), in plain black.

For both cvs, final SY was tightly related to seed numbers (*r^2^* ≥ 0.856; *P<0.001*) and to a much less extent to individual seed weight (*r^2^* ≤ 0.086; *0.01<P<0.10*).

When environments were sorted in four groups depending upon the combination of mean temperature and relative water balance (RWB) along the cycle (Fig. 2E), it could be observed that most part of the experiments (13 cases) fell within the warm and dry category (i.e. RWB≤0 and mean temperature ≥22°C), followed by the cool and dry (7 cases), then the warm and wet (5 cases) and finally the cool and wet (2 cases). Considering dry environments (i.e. RWB≤0), the mean relative SY (RSY) was +8.6% (i.e. TG > WT). Within this group, the subgroup warm and dry had a mean RSY of +10.5%, whereas the dry and cool subgroup had a mean RSY of +5.1%. The mean RSY of wet environments was +3.6%, being +5.2% for the warm and wet and almost null (−0.5%) for the cool and wet. Although the wet and cool environmental conditions are not preponderant among common growing conditions experienced by soybean crops, further evaluation is necessary in order to test the TG efficacy under such condition. It is important to highlight that cases with negative RSYs were scattered across all environmental categories. Therefore, negative values could not be attributed to a specific environmental condition but to some other factor/s that caused a detriment to the presence of *HaHB4.*

### Differential traits between transgenic and WT soybean grown in the greenhouse

Aiming at understanding which physiological traits could be responsible for the drought/warm tolerant phenotype observed in *HaHB4*-transgenic plants, a morpho-physiological evaluation was performed on plants grown in the greenhouse under well-watered or water-deficit conditions (Fig. 3A). Transgenic b10H plants exhibited a trend to increased SY in both conditions, even though SY of both genotypes was significantly affected (*P<0.05*) by water deficit (Fig. 3B). Differences in SY were accompanied by similar trends in seed numbers but not in seed weight (Fig. 3B). Cumulative water use during the water-deficit conditions, normalized by cumulative water use by well-watered plants, was mainly a consequence of the enhanced water use by TG b10H under well-watered conditions (Fig. 3C). Transgenic plants tended to be shorter and to have more branches, internodes and pods per plant than the control W82 (Fig. 3D).

**Figure 3.**
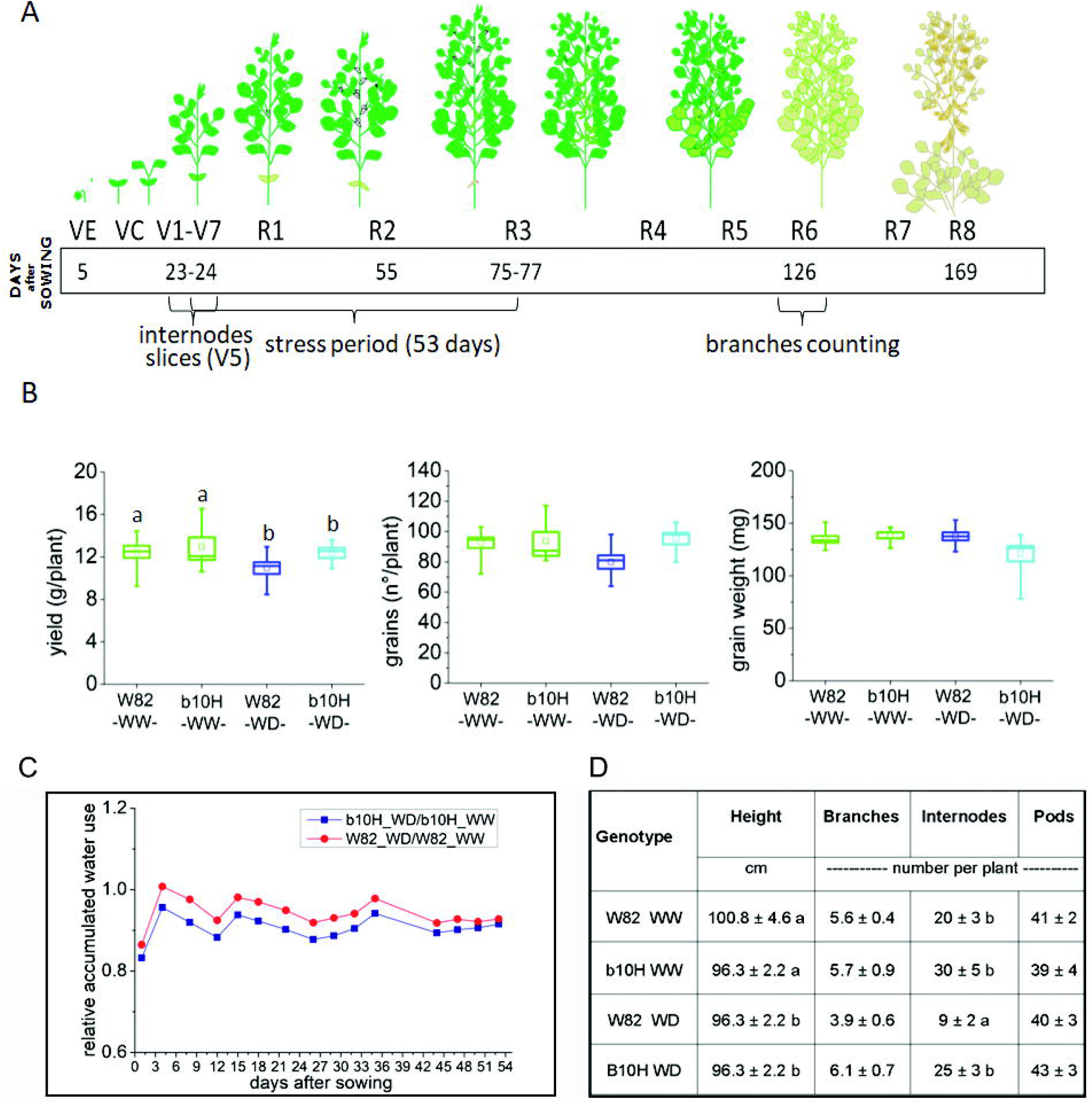
Transgenic HaHB4 soybean grown in the greenhouse under contrasting water regimes differs from its control W82 in seed yield and its components. (A) Schematic representation of soybean developmental stages and dates in which drought stress treatment was applied as well as dates in which data were collected. (B) Comparison of seed yield and yield components (seed number and individual seed weight) per plant in well-watered (WW) and water-deficit (WD) plants; water deficit was applied in the period indicated in A. Different letters mean significant differences between samples (*P* < 0.05). (C) Relative cumulated water use during the water-deficit period. Numbers (as %) illustrate water volume used by b10H (relative to W82 considered as 100%) in the more representative points during WD and WW treatments. (D) Comparison of plant height as well as the number of branches, internodes and pods among treatments described in (B). In (D), data are mean ± standard errors, and means followed by the same letter within a column do not differ at *P<0.05.*

Diameter of epicotyls, and first, second and third internodes were measured, indicating that those of b10H tended to be wider than those of W82 in stressed plants (Fig. 4A). The same trend can be observed for xylem area whereas all these differential characteristics were less remarkable in well-watered plants (Figs. 4C-F and Supplementary Fig. 2).

**Figure 4.**
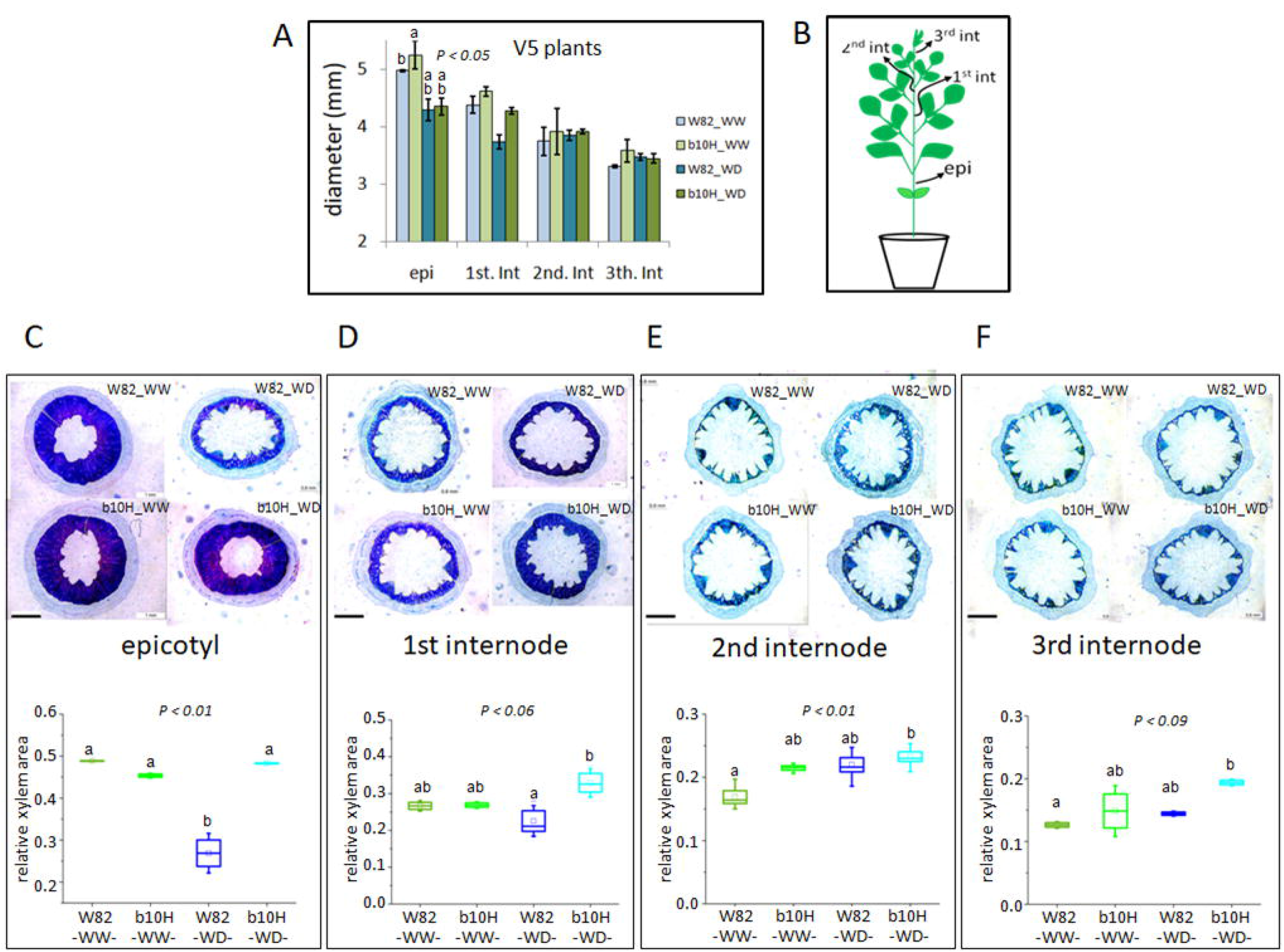
HaHB4 soybean plants exhibit wider stem diameter and larger xylem area than controls. (A) Stem diameter of V5 plants in well-watered (WW) and water-deficit (WD) plants. (B) Schematic representation of a soybean plant indicating the stem sections analyzed in (C). (C-F) Histological epicotyl, hypocotyl and stem cuts stained with safranine fast green and quantification of xylem area (lower panel). Xylem area was highlighted in cross stem sections (upper panel) and xylem relative areas (bottom panel) are shown. Bar length = 1mm. Different letters mean significant differences according to Tukey comparisons with the indicated *P* values.

### Plant phenotyping in field trials indicates significant differences between transgenic b10H and controls

Aiming at knowing whether the differential architectural and physiological traits observed in the greenhouse were conserved in plants grown in the field, production traits plus several physiological traits were assessed in TG b10H and WT W82 soybeans in experiments performed at the IAL site during 2017-2018 (Fig. 5A). Plants were irrigated but experienced some degree of above-optimum temperatures (i.e. heat stress) along the cycle (Supplementary Table 1). No significant differences were detected in evaluated traits between genotypes at R3. By contrast, b10H plants had a significantly (*P<0.05*) higher photosynthesis rate than W82 at R5 and R6 (Fig. 5B), a trend also observed for light interception during seed filling (R6) and crop biomass (Fig. 5C). Differences in crop biomass were accompanied by significantly (*P<0.05*) increased SY (Fig. 5C) and seed numbers (Fig. 5C). Increased seed numbers overcompensated the reduction registered in individual seed weight (Fig. 5C). Differences in seed numbers were driven by the augmented number of branches and pods registered for b10H as compared to W82 plants (Figs. 5D and 5E). Finally, and similar to the phenotype observed in the greenhouse experiment, hypocotyl diameter and xylem area were larger in b10H than in W82 (Fig. 6 and Supplementary Fig. 3).

**Figure 5.**
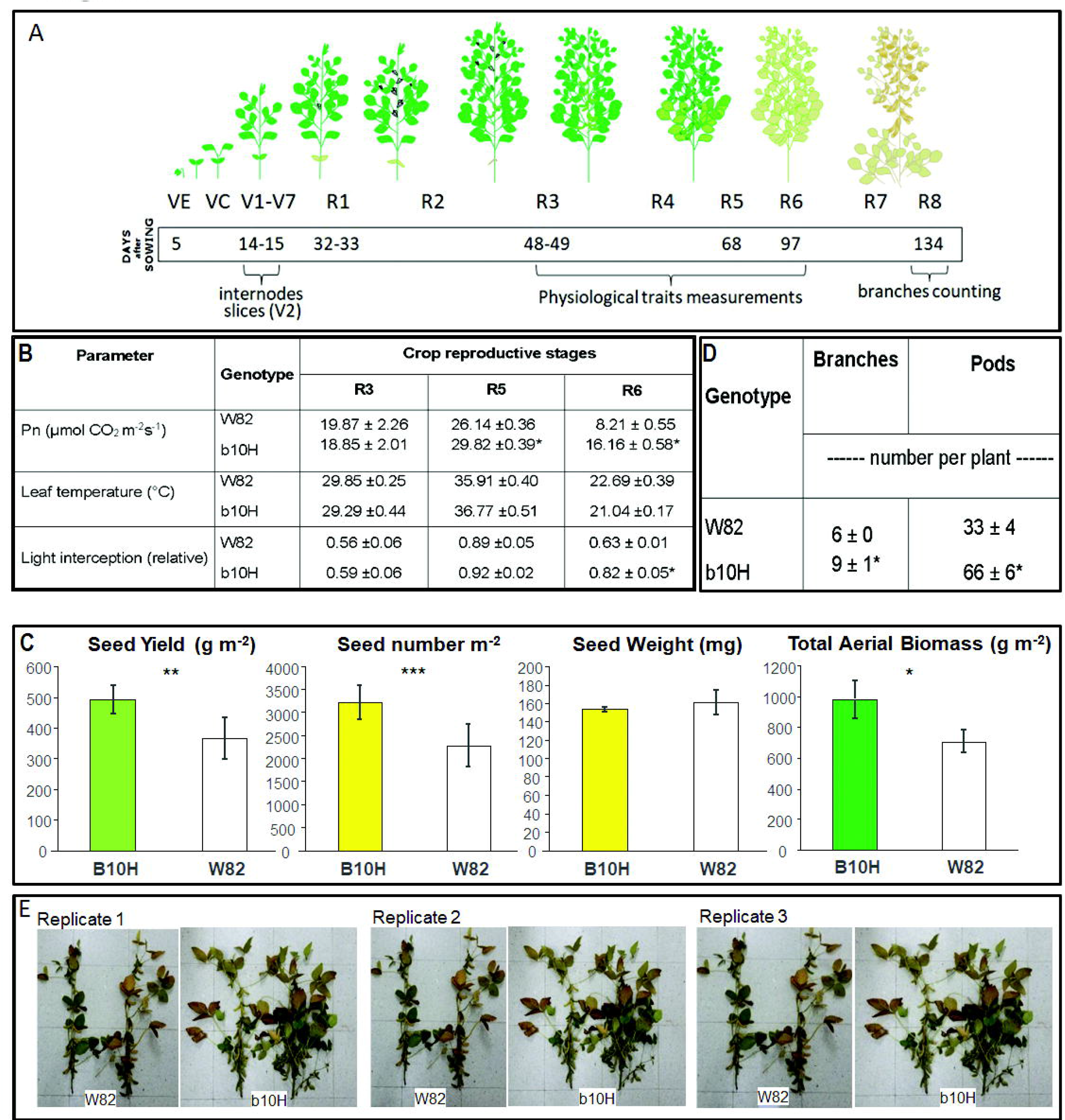
Seed yield components differ between field-grown transgenic b10H and wild type W82 in the warm-wet environment of the the IAL site. (A) Details of assessed characteristics and dates of data collection during soybean crop cycle in a field trial carried out at the IAL site. Comparison between b10H and W82 of (B) physiological traits measured at different growth stages (Pn: net photosynthesis), (C) seed yield and yield components (seeds number, seed weight and total aerial biomass), and (D) number of branches and pods per plant. (E) Illustrative pictures of plants collected from all replicates. In (B) and (D), data are the mean ± standard errors and asterisks indicate significant differences between b10H and W82 at *P* = 0.05. In (C), error bars represent SEM×2 and asterisks indicate significant differences between b10H and W82 (*,**,*** for *P* values of 0.10, 0.05 and 0.01, respectively)

**Figure 6.**
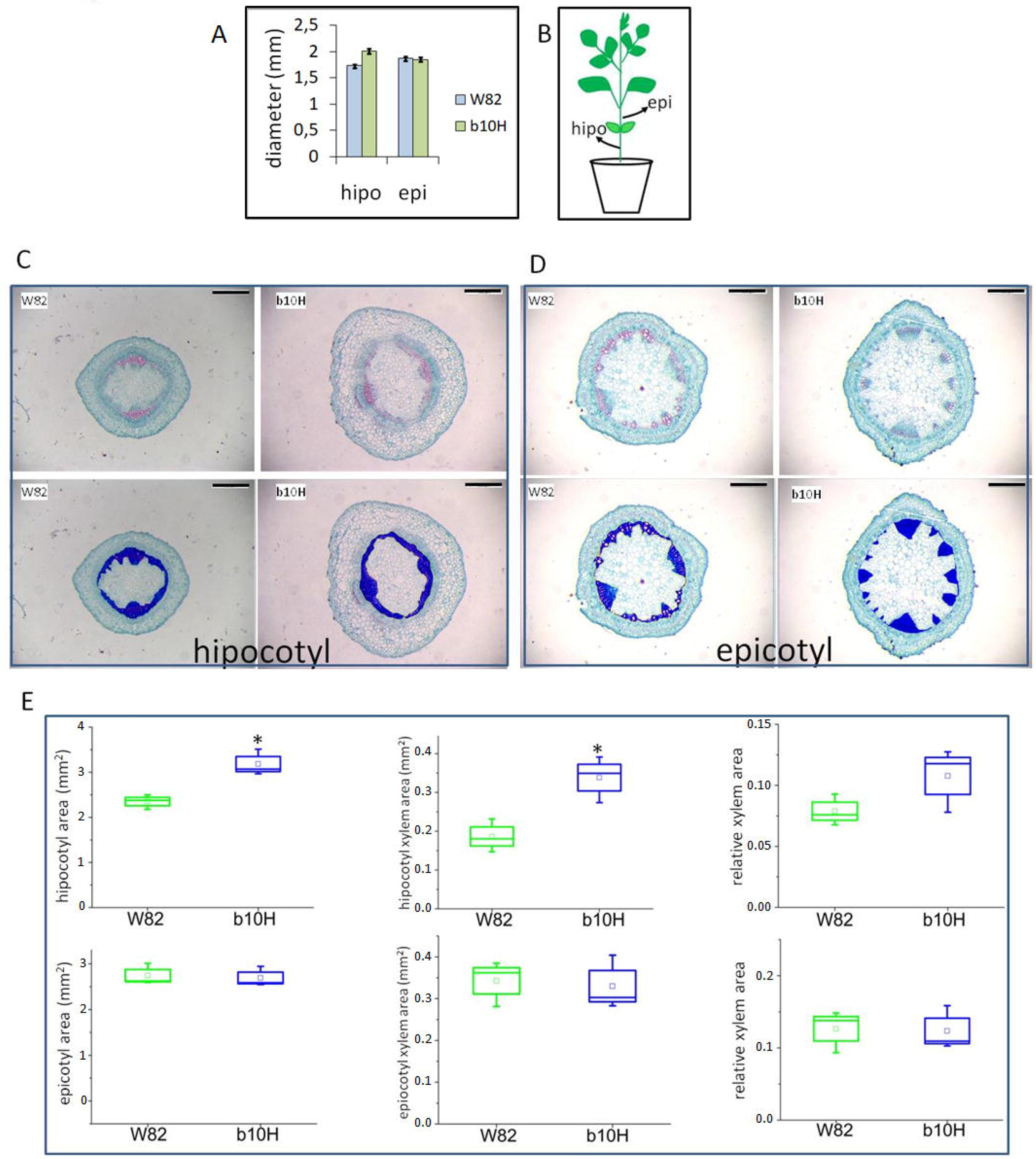
Architectural characteristics differ between transgenic b10H and wild type W82 in plants grown in the IAL site. (A) Internodes width (hypocotyl, epicotyl) in V2 plants. (B) Illustration of a plant indicating which tissues were harvested for morphological analyses. (C-D) Illustrative pictures with standard (upper panel) or highlighted (bottom panel) xylem area of cross stem sections obtained after safranine-fast green staining. (E) hipocotyl (upper panel) and epicotyl (bottom panel) cross stem section areas. Total internodes and xylem area, and xylem relative area were plotted and calculated using three biological replicates. Bar length = 0.5 mm. Different letters means significant differences according to Tukey comparisons with the indicated *P* values.

Also during 2017-2018, when summer crops in the temperate region of Argentina were exposed to a severe drought caused by a *La Niña* phase of the *ENSO* phenomenon, a field-based analysis of SY determination under two contrasting water regimes (WD: water deficit; WW: well-watered) was performed at the Pergamino site. Rainfall exclusion plus differential irrigation produced a large contrast in total crop evapotranspiration (ET_C_) between WD and WW plots (Fig. 7A). Soil water survey included the topmost 185 cm and was performed from sowing+23 d to R7. In both conditions, water use of the TG cv. b10H was higher (*P<0.05*) than water use computed for the WT cv. W82. (17.3% in WD and 27.2% in WW). The physiological analysis indicated that the former outyielded the latter under WD (43.4%) with no penalization under WW (Fig. 7B). As observed in the IAL experiment, increased SY registered in b10H respect to W82 under drought was driven by increased crop (44.5%; Fig. 7C) and pod biomass (52.6%; Fig.7D) as well as by increased pod (73.3%; Fig. 7G) and seed numbers (78.9%; Fig. 7H). Drought produced no significant difference between cvs in biomass partitioning to pods (Fig. 7E) or to seeds (Fig. 7F), whereas individual seed weight in this condition was larger for W82 than for b10H (24.6%; Fig. 7I). Based on trends registered for crop water use (ETC) and production traits, a remarkably higher (≥22%) water use efficiency (WUE) was computed for TG than for WT cultivars exposed to drought. This trend held for biomass (WUE_B,ETc_ of 2.3 g m^−2^mm^−1^ for the transgenic and of 1.9 g m^−2^ mm^−1^ for W82) as well as for seed WUE (WUE_SY,ETc_ of 0.91 g m^−2^ mm^−1^ for the transgenic and of 0.74 g m^−2^ mm^−1^ for W82).

**Figure 7.**
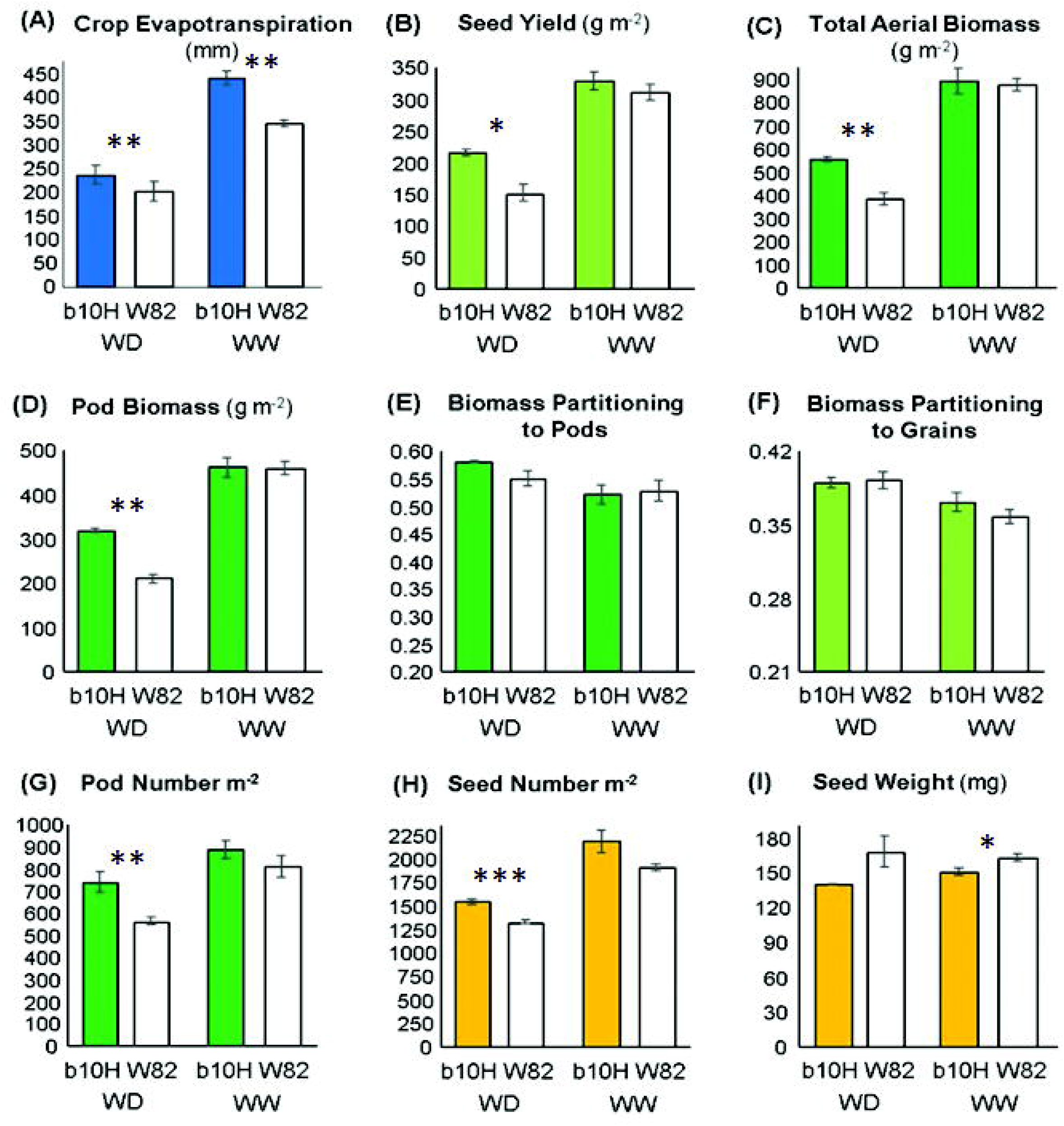
Field-based physiological analysis of seed yield determination under contrasting water regimes. Mean values of different traits evaluated at harvest for transgenic cv. b10H and WT cv. W82 grown under water-deficit (WD) and well-watered (WW) conditions at the Pergamino site. (A) Crop evapotranspiration during the cycle. (B) Seed yield. (C) Total aerial biomass. (D) Pod biomass. (E) Biomass partitioning to pods (ratio between pods biomass and total aerial biomass). (F) Biomass partitioning to seeds or harvest index (ratio between seed yield and total aerial biomass). (G) Pod numbers. (H) Seed numbers. (I) Individual seed weight. Error bars represent SEM × 2 and asterisks indicate significant differences between b10H and W82 (*,**,*** for *P* value of 0.10, 0.05 and 0.01, respectively).

### Different molecular pathways are altered in transgenic soybean expressing HaHB4

Using RNA-Seq of TG versus WT plants, we identified 743 differentially expressed genes (DEGs, FDR adjusted p-value < 0.05, Fig. 8A, Supplementary Table 2), of which 120 presented an absolute log2-fold change greater than one. An inspection of the DEGs based on potential orthologous genes from Arabidopsis showed that there were genes previously associated to the heterologous expression of *HaHB4*, as lipoxygenases, trypsin inhibitors (Manavella *el al.*, 2008) and the Cu/Zn superoxide dismutase CSD1 (Manavella *el al.*, 2006). There were also many DEGs related to heat, as the homologues of heat shock protein genes AT-HSC70-1 (AT5G02500), AT-HSFB2A (AT5G62020), Hsp81.4 (AT5G56000), and the homologue of the heat related gene *HOT5* (AT5G43940, also known as *GSNOR*, Lee *et al.*, 2008).

**Figure 8.**
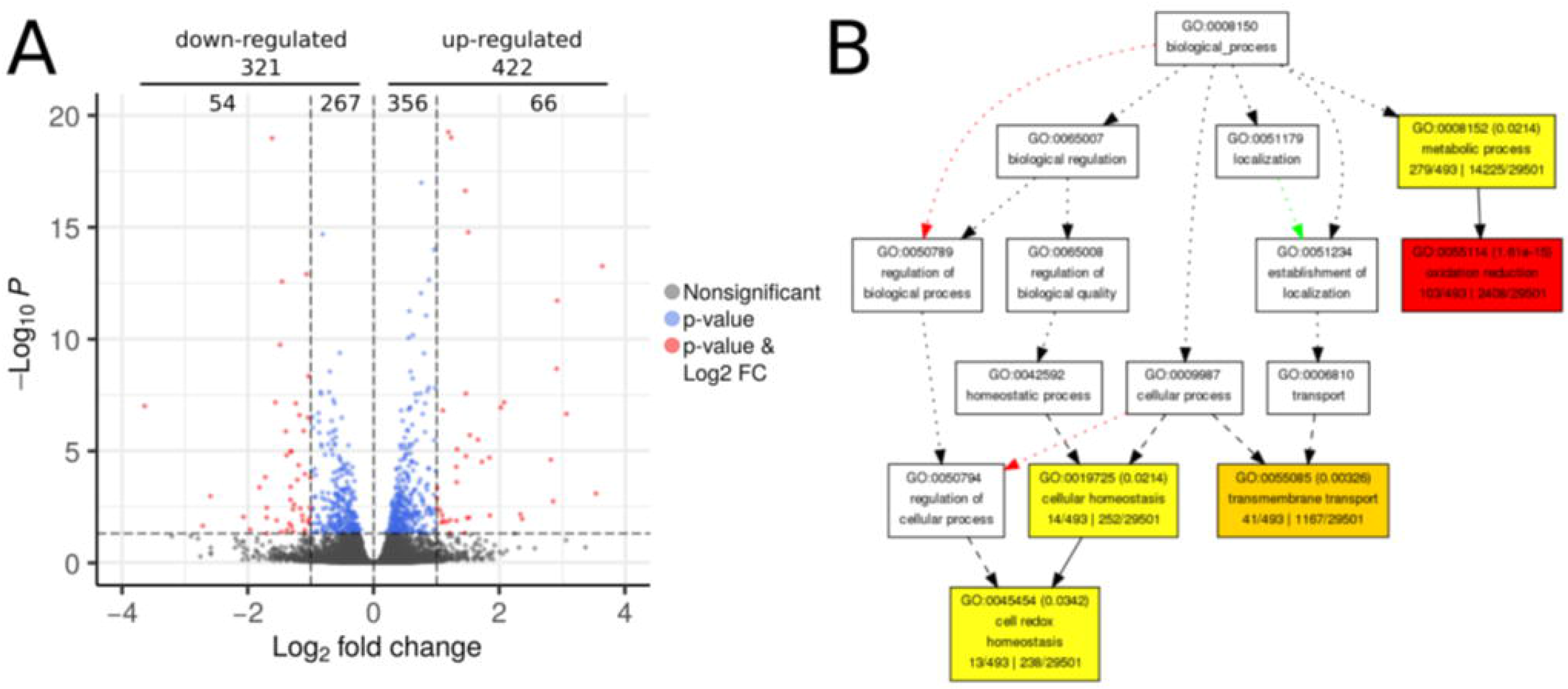
Differentially expressed genes in soybean HaHB4 compared to W82 in field trials. (A) Volcano plot of differentially expressed genes (DEGs) determined with RNA-seq in transgenic versus W82 plants. DEGs had an FDR adjusted p-value below 0.05 as indicated (horizontal cutoff). Additionally, genes above or below an absolute log2-fold change of one were differentially colored and their number is in the upper margin of the plot. A few genes were omitted as their absolute fold change was greater than the chosen axis graph limits. (B) Directed acyclic graph of GO terms including enriched categories of the BP (biological process) type.

A gene ontology enrichment analysis on all DEGs revealed “oxidation reduction”, “cell redox homeostasis” and “transmembrane transport” as interesting significantly enriched BP terms (*P < 0.05*, Fig. 8B, Supplementary Table 3). Among MF terms (Supplementary Table 3), some more descriptive categories were enriched, such as “protein disulfide oxidoreductase activity”, “iron ion binding”, “metal ion binding” and “peroxidase activity” (Supplementary Table 3).

## Discussion

Second generation of transgenic crops (i.e. those aimed to abiotic stress tolerance) did not reach the market yet, with the sole exception of drought-tolerant maize (http://www.isaaa.org/gmapprovaldatabase/) transformed with the bacterial RNA chaperons CspB and CspA (Castiglioni *et al.*, 2008). Besides the difficult and long regulatory processes that transgenic crops must go through, an additional constraint for this second generation is the non-universal nature of abiotic stresses, a characteristic that contrasts with the qualitative nature of biotic-oriented TG crops like the emblematic RR soybeans and Bt maize. Mentioned constraint applies particularly to drought, which may display in a broad spectrum of alternatives derived from multiple combinations of growth stages, intensities and durations along the cycle (Chapman *et al.*, 2000; Tardieu, 2012).

Although the vast literature about drought tolerant transgenic plants, mostly demonstrated in models and in controlled conditions (Skirycz *et al.*, 2011; Passioura 2012), it is possible that the huge investments required to release drought-tolerant crops were aborted at different stages. Hence, the lack of drought tolerant crops in the market is likely caused by experimental failures experienced when technologies tested in model plants and controlled conditions were surveyed in field-grown crops.

HaHB4 is a sunflower transcription factor whose expression is up-regulated by water deficit (Gago *et al.*, 2002). Its ectopic expression in Arabidopsis leads to drought tolerant plants following complex physiological mechanisms that do not include stomata closure, a typical plant response to deal with water deficit involving a decrease in ethylene sensitivity (Dezar *et al.*, 2005; Manavella *et al.*, 2006). It was recently demonstrated that *HaHB4* was able to confer drought tolerance to wheat in field trials (Gonzalez *et al.*, 2019) although the evolutionary long distance between sunflower, Arabidopsis and wheat. In this work we demonstrated that soybeans transformed with *HaHB4* also performed better than its WT control in stress-prone field conditions, particularly in warm and dry environments. These results were especially interesting because the expected drought-tolerant phenotype observed in Arabidopsis and wheat expanded to a drought/heat-tolerant one in the case of soybeans, which is a promising outcome for future climatic scenarios (IPCC, 2014).

Many independent events were obtained at the beginning of the research, including several ones with the combination of the *HaHB4* own promoter and the first intron of the Arabidopsis *COX5c* gene acting as an enhancer (Curi *et al.*, 2005; Cabello *et al.*, 2007). However, after several field trials in different environments, the most robust events were a5H and a11H (HaHB4 expression driven by the *35S CaMV* promoter) and b10H (expression driven by the *HaHB4* promoter). Such events tended to outyield the WT parental cv in all trials. Among them, b10H was the best one and further studies were carried out only with this event.

Seed yield variation considering water regimes and temperatures strongly suggested that the best target environments for transgenic b10H soybeans are the warm and dry ones, in which it clearly outyielded controls (Fig. 2). Notably, the best performances were obtained in the droughted experiment of Pergamino and the irrigated experiment of the IAL sites, both developed during the *La Niña* phase of the ENSO phenomenon that took place during 2017-2018 (https://origin.cpc.ncep.noaa.gov), which brought below normal rainfall together with episodes of above-optimum temperatures during the cycle of summer crops in the Pampas region of Argentina (Supplementary Table 1).

In all cases in which TG outyielded controls, the response was associated to improved seed numbers that overcompensated the decline registered in individual seed weight. These characteristics (i.e. partial trade-off between seed yield components) were also observed in the greenhouse experiment. Collectively, results highlight the importance of improved crop growth (i.e. resource acquisition and allocation to reproductive organs) during the critical period for seed establishment (Vega et al., 2001) as well as of the necessary improvement in the crop photosynthetic capacity during seed filling (Borrás et al., 2004) to avoid the mentioned trade-off between seed yield components. . Soybean SY history of the past 90 years has been recently revised, and potential targets to achieve yield improvement were proposed (Ainsworth *et al.*, 2012). As for cereals (Slafer *et al.*, 2015), optimization of carbon utilization/delivery to avoid flower abortion (i.e. improved fruiting efficiency) is among such targets in soybeans (Egli and Bruening, 2002; Kantolic and Slafer, 2005), for which Ainsworth *et al.* (2012) proposed to advance molecular breeding techniques aimed to the regulation of flower initiation and abortion. In this sense, TG b10H plants seem a promising genetic resource for future studies.

An outstanding feature of TG b10H plants was their enhanced water use under well-watered conditions (Figs. 3C and 7A), particularly in field-grown plots. Because no evident difference was registered in the phenotype of WT and TG plants in this condition (i.e. identical visual canopy characteristics), differences cannot be ascribed to a contrasting participation of soil evaporation in total crop water use and can only be linked to enhanced transpiration of the TG genotype. This trend may be ascribed to the enhanced xylem area and stem diameter observed in the TG phenotype, traits that may contribute to increase hydraulic conductivity and water use by crops (Richards and Passioura, 1981) and have been recently associated with increased yield of Arabidopsis as well as of sunflower plants (Cabello and Chan, 2019). However, differences in water uptake between TG and WT cvs declined markedly (Fig. 7A) or almost disappeared (Fig. 3C) under water deficit, suggesting that described benefits observed in well-watered environments may have been partially or totally compensated in response to drought, probably by enhanced stomata controlled of gas exchange in b10H respect to W82 (Sadock and Sinclair, 2009). Nevertheless, such control may have had a larger effect on water loss than on CO_2_ fixation (Liu *et al.*, 2005), a response supported by the pronounced increased in WUE registered under drought for b10H compared with W82. Such response was not biased by differences related to the soil component of crop evapotranspiration, which was minimized in this growing condition. Collectively, described results are in good agreement with the slow-wilting soybean phenotype characterized by Fletcher *et al.* (2007), which might allow water conservation in drought conditions with no yield penalization in potential environments (Devi *et al.*, 2014).

The acceleration of senescence by water stress during seed filling has been documented in soybean (de Souza *et al.*, 1997; Brevedan and Egli, 2003). Therefore, the delay in senescence reported here would be expected if the transgene promotes a reduced sensitivity to ethylene as reported in Arabidopsis (Manavella *et al.*, 2006). Delayed senescence matched the delayed in phenology registered only for the R7 stage (i.e. late in the cycle), which could be visually assessed in several but not all field trials. The more surprising response observed in TG soybean was the tolerance to warm/dry growing conditions, which underscores the target environments for this technology. Such response was not registered in Arabidopsis-HaHB4 neither in field-tested wheat-HaHB4. In the case of the former, because model plants analyzed for drought tolerance were always grown under controlled temperature (Dezar *et al.*, 2005; Manavella *et al.*, 2006, 2008) but were never exposed to above-optimum ones. In the case of the latter, because the winter crop did not experience the combined effect of drought and high temperature episodes along the cycle, except during grain filling of a few experiments (González *et al.*, 2019). Further investigations will be necessary to elucidate whether this behavior (warm/dry tolerance) is universal to all HaHB4-bearing species (i.e. gene specific) or it is limited to soybean (i.e. HaHB4 × species interaction).

Regarding the transcriptome of transgenic soybean, it is tempting to speculate that conserved mechanisms are displayed in different species even when they must be corroborated to support this hypothesis. This is because despite the great difference between 3-week-old culture chamber grown Arabidopsis plants (Manavella *et al.*, 2006, 2008) and R5 soybeans grown in the field, it is remarkable to observe that non-typically drought-responsive genes were differentially regulated and several encoding lipoxygenaes, trypsin inhibitors and Cu/Zn superoxide dismutase appeared as regulated in TG plants of both species. The surprise was to find heat shock related genes differentially regulated in soybean like homologues of AT-HSC70-1, AT-HSFB2A, Hsp81.4 and HOT5 (Lee *et al.*, 2008), which support the mentioned tolerance to high temperatures registered in current research and will be investigated in the near future. Moreover, the GO term “cell redox homeostasis”, known to be important under many stressful conditions (Vinocur and Altman, 2005), was enriched among DEGs. Experiments will be aimed at defining if such regulation persists under other environmental conditions or it is displayed by HaHB4 only when plants are subjected to warm/dry environments.

Finally, soybean commercial varieties derived from the b10H event are currently being developed by multiple technology licensees. The event (named IND-ØØ41Ø-5 for regulatory and commercial release) has been conditionally approved for commercialization in Argentina in 2015 (https://www.argentina.gob.ar/agroindustria/alimentos-y-bioeconomia/ogm-comerciales), subject to Chinese importation clearance for food and feed use (according to feed safety assessment principles; Parrott *et al.*, 2010), which is still pending. Brazil (https://cib.org.br/produtos-aprovados/) and more recently the United States (https://www.aphis.usda.gov/aphis/ourfocus/biotechnology/permits-notifications-petitions/petitions/petition-status) have approved also the event for production and consumption purposes. Together, these three countries represent about 80% of the global soybean production. Elite varieties are currently being multiplied and a few thousand hectares are expected to go into production in the coming crop cycle in the southern Hemisphere. The technology is expected to be broadly launched in South America in 2020-21, under the HB4 brand.

## Supplementary material

**Supplementary Table 1.** General growing conditions along the cycle experienced by soybean crops in 27 experiments

**Supplementary Table 2.** Differentially expressed genes between transgenic and W82 control plants

**Supplementary Table 3.** GO Terms analysis of 743 differentially expressed genes

**Supplementary Figure 1.** Response of grain yield and grain yield components of W82 and b10H to their corresponding environmental indexes

**Supplementary Figure 2.** Illustrative images of histological stem cuts of W82 and HaHB4 transgenic plants

**Supplementary Figure 3.** Illustrative images of histological stem cuts of W82 and b10H plants in the IAL site.

## Acknowledgments

This work was supported by Agencia Nacional de Promoción Científica y Tecnológica, PICT 2015 2671. Field trials shown in Figures 1 and 2 were supported by INDEAR (ANR ANPCyT-FONTAR 209/09 and 295/13).

In this work computational resources from the PIRAYU cluster, acquired with funds from project AC-00010-18, Res.No117/14, from ASACTEI.

We thank Dr. Patricia Miranda for critical reading of this manuscript.

## Authors contributions

SAB, MEO, KFR and JVC carried out physiological and morphological evaluations at Pergamino and IAL sites; KFR prepared RNA and evaluated transcript levels; KFR and ALA did transcriptome analyses; CA and MP performed IRGA analyses; MC, GW and FT designed and carried out field trials. FT, MP, MEO and RLC conceived, designed and wrote the manuscript.

