## Supplemental Fig 1 for "Successful field performance in dry-warm environments of soybean expressing the sunflower transcription factor HaHB4"

**Supplementary Figure 1.**

Comparison between the transgenic cv. b10H and the parental cv. W82 of seed yield (A), seed number (B), and individual seed weight (C) across all evaluated environments (Supplementary Table 1). The mean value of each environment corresponds to the average of all tested genotypes and for each trait is described as an environmental index (EI) . Fitted models in (A) differed at $P<0.05$ and indicated that b10H will outyield W82 across all environments with seed yield lower than 4898 kg ha$^{-1}$, a threshold never met in current research. No cross-over interaction was detected for models fitted to seed numbers (B) and seed weight (D). Ordinates of models fitted in (B) and (C) differed at $P<0.0001$.

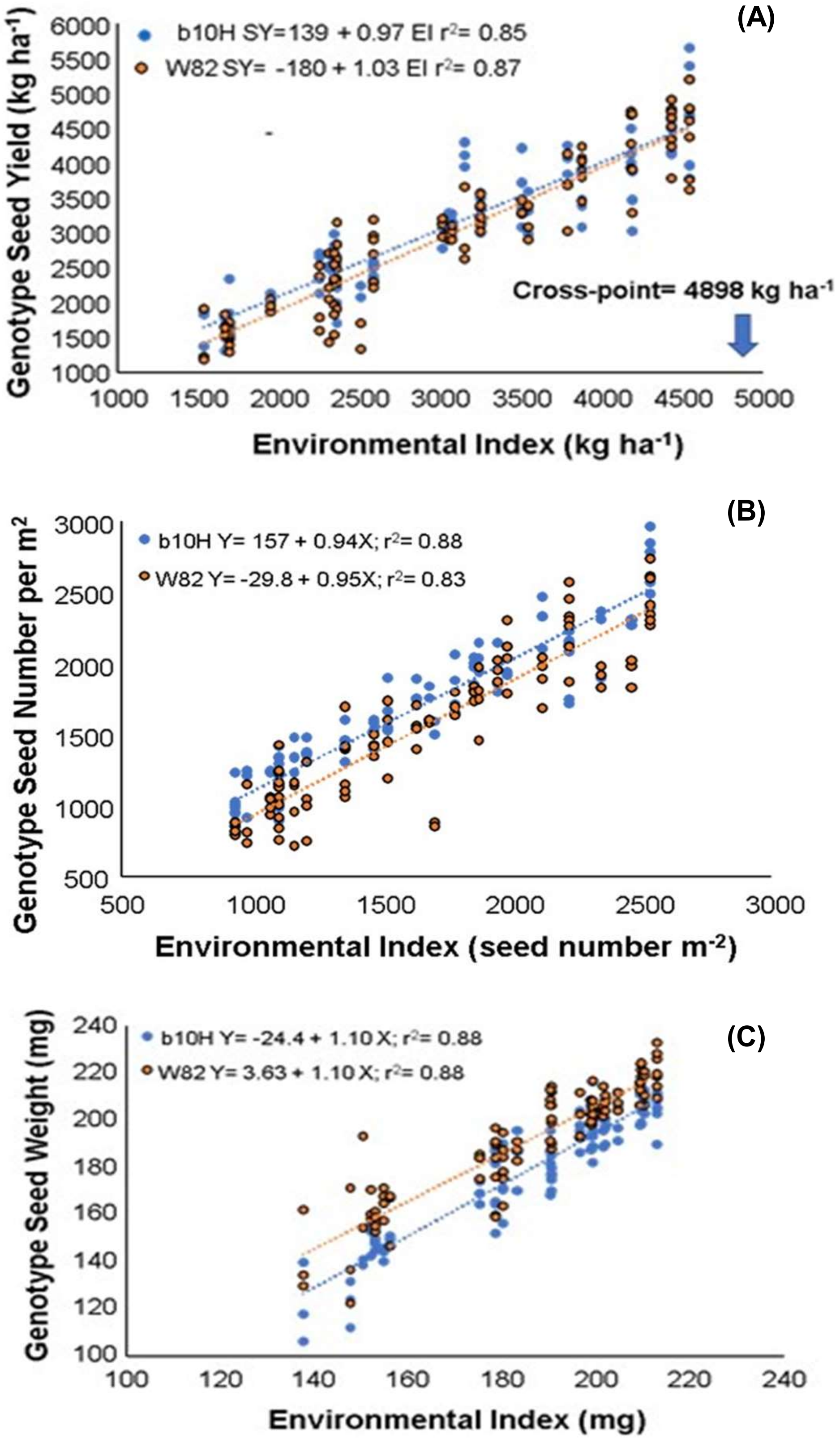
