## Supplementary figures and images for "Successful field performance in dry-warm environments of soybean expressing the sunflower transcription factor HaHB4"

### Supplemental Fig2

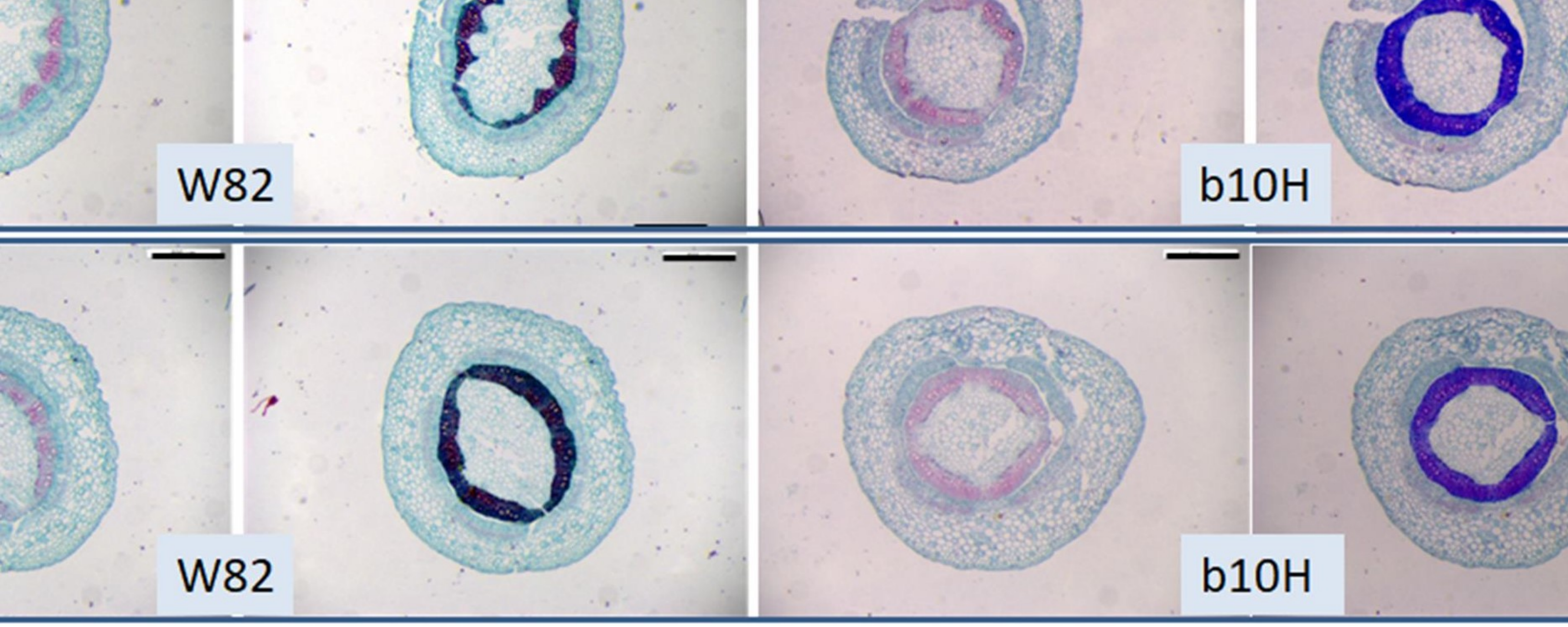

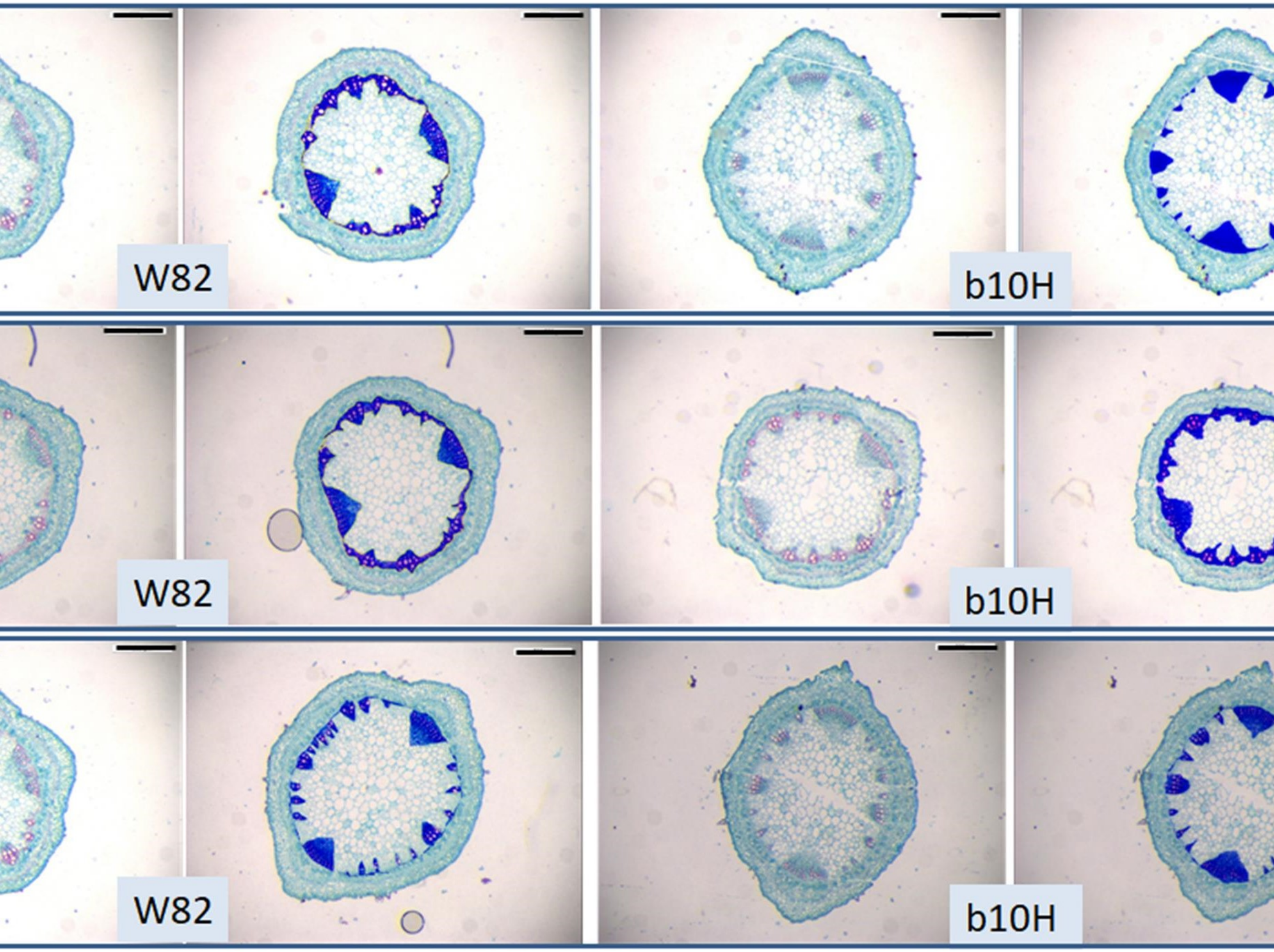
