## Supplemental Table 1 for "Successful field performance in dry-warm environments of soybean expressing the sunflower transcription factor HaHB4"

**Supplementary Table 1. Description of experiments.**

Data correspond to weather coditions experienced during the whole cycle by field-grown soybean crops in 27 experiments performed across 6 years and 14 sites of Argentina. Sites are indicated in the map of Fig. 2A. Different water regimes (IR: irrigated; R: rainfed; WD: water deficit with rain-out shelters), sowing dates (ES: early sowing; DS: delayed sowing), and phosphorus fertilizer rates (MAP: monoammonium phosphate, in kg $ha^{-1}$) were included in the network. Groups refer to the scope of differente experiments (G1 for event selection, G2 for genotype × environment evaluation and G3 for physiological analysis). Within each column, the color scale goes from dark green for the smallest value to dark red for the largest value, except for Rain+IR (opposite scale). ID: identification; SRc: cumulative incident solar radiation; Tmax: mean daily maximum temperature; Tmin: mean daily minimum temperature; Tmean: mean daily mean temperature; PET: potential evapotranspiration; Rain+IR: rainfall+irrigation; WB: water balance (WB= Rain+IR-PET)); RWB: relative water balance (RWB= $\frac{\text{Rain+IR}-\text{PET}}{\text{PET}}$).

| Site in map | Harvest year | Experiment description | Water management | ID | Groups | SRc | Tmax | Tmin | Tmean | PET | Rain+IR | WB | RWB |
|---|---|---|---|---|---|---|---|---|---|---|---|---|---|
| | | | | | | MJ $m^{-2}$ | °C | °C | °C | mm | mm | mm | |
| 1 | 2013 | 01. Aranguren ES | Rainfed | AR1a | G1, G2 | 2726 | 30.8 | 17.2 | 24.0 | 618 | 365 | -253 | -0.41 |
| 1 | 2013 | 02. Aranguren DS | Rainfed | AR1b | G1, G2 | 2263 | 30.2 | 16.3 | 23.1 | 641 | 310 | -331 | -0.52 |
| 1 | 2014 | 03. Aranguren MAP_0 | Rainfed | AR2a | G2 | 1714 | 29.1 | 17.7 | 23.4 | 360 | 325 | -35 | -0.10 |
| 1 | 2014 | 04. Aranguren MAP_100 | Rainfed | AR2b | G2 | 1714 | 29.1 | 17.7 | 23.4 | 360 | 325 | -35 | -0.10 |
| 2 | 2013 | 05. Carmen de Areco ES | Rainfed | CA1 | G1, G2 | 2990 | 29.8 | 15.6 | 22.8 | 644 | 496 | -148 | -0.23 |
| 2 | 2013 | 06. Carmen de Areco DS | Rainfed | CA2 | G1, G2 | 2915 | 27.5 | 13.9 | 20.6 | 615 | 379 | -236 | -0.38 |
| 3 | 2013 | 07. Corral de Bustos | Rainfed | CB | G1, G2 | 2765 | 29.0 | 15.1 | 21.8 | 574 | 337 | -237 | -0.41 |
| 4 | 2013 | 08. Chilibroste | Rainfed | CH | G1, G2 | 2908 | 30.6 | 15.7 | 22.8 | 646 | 407 | -239 | -0.37 |
| 5 | 2012 | 09. Hughes | Rainfed | HU1 | G1, G2 | 2571 | 25.8 | 13.5 | 19.3 | 766 | 539 | 228 | 0.30 |
| 5 | 2013 | 10. Hughes | Rainfed | HU2 | G1, G2 | 2668 | 28.0 | 14.8 | 21.3 | 575 | 341 | -234 | -0.41 |
| 6 | 2018 | 11. IAL-Santa Fe | Irrigated | IAL | G2, G3 | 2884 | 32.4 | 19.1 | 25.6 | 688 | 766 | 78 | 0.11 |
| 7 | 2013 | 12. Landeta | Rainfed | LA | G1, G2 | 2272 | 28.2 | 14.3 | 21.0 | 489 | 329 | -160 | -0.33 |
| 8 | 2010 | 13. Liborio Luna R | Rainfed | LL1 | G2, G3 | 2586 | 29.0 | 14.6 | 21.4 | 564 | 404 | -161 | -0.28 |
| 8 | 2010 | 14. Liborio Luna IR | Irrigated | LL2 | G2, G3 | 2586 | 29.0 | 14.6 | 21.4 | 564 | 570 | 5 | 0.01 |
| 9 | 2013 | 15. Monte Buey | Rainfed | MB1 | G1, G2 | 3440 | 28.9 | 15.0 | 21.7 | 744 | 323 | -421 | -0.57 |
| 9 | 2014 | 16. Monte Buey | Rainfed | MB2 | G2 | 2194 | 28.2 | 15.6 | 21.8 | 478 | 580 | 102 | 0.21 |
| 10 | 2018 | 17. Pergamino WD | Water Deficit | Pe1_D | G2, G3 | 2637 | 31.6 | 16.5 | 23.9 | 602 | 152 | -451 | -0.75 |
| 10 | 2018 | 18. Pergamino IR | Irrigated | Pe1_I | G2, G3 | 2637 | 31.6 | 16.5 | 23.9 | 602 | 332 | -270 | -0.45 |
| 10 | 2019 | 19. Pergamino IR | Irrigated | Pe2_I | G2, G3 | 2624 | 27.8 | 16.2 | 22.1 | 569 | 682 | 113 | 0.20 |
| 10 | 2019 | 20. Pergamino R | Rainfed | Pe2_R | G2, G3 | 2624 | 27.8 | 16.2 | 22.1 | 569 | 806 | 237 | 0.42 |
| 11 | 2010 | 21. Quimili | Rainfed | QU | G1, G2 | 1868 | 32.5 | 20.3 | 26.1 | 451 | 377 | -75 | -0.17 |
| 12 | 2014 | 22. Roldán ES | Rainfed | RO1 | G2 | 2169 | 30.3 | 18.3 | 24.2 | 502 | 475 | -27 | -0.05 |
| 12 | 2014 | 23. Roldán DS | Rainfed | RO2 | G2 | 2230 | 27.7 | 16.2 | 21.8 | 480 | 571 | 91 | 0.19 |
| 13 | 2013 | 24. San Agustín ES | Rainfed | SA1 | G1, G2 | 3172 | 25.6 | 12.0 | 18.4 | 637 | 350 | -287 | -0.45 |
| 13 | 2013 | 25. San Agustín DS | Rainfed | SA2 | G1, G2 | 3002 | 25.5 | 11.9 | 18.3 | 602 | 376 | -226 | -0.38 |
| 14 | 2013 | 26. Villa Saboya | Rainfed | VS1 | G1, G2 | 2426 | 29.0 | 14.7 | 21.6 | 528 | 252 | -276 | -0.52 |
| 14 | 2014 | 27. Villa Saboya | Rainfed | VS2 | G2 | 2306 | 28.9 | 15.7 | 22.2 | 508 | 459 | -49 | -0.10 |
